# Deep neural net tracking of human pluripotent stem cells reveals intrinsic behaviors directing morphogenesis

**DOI:** 10.1101/2020.09.21.307470

**Authors:** David A. Joy, Ashley R. G. Libby, Todd C. McDevitt

**Affiliations:** UC Berkeley-UC San Francisco Graduate Program in Bioengineering, San Francisco; Gladstone Institutes, San Francisco; Developmental and Stem Cell Biology PhD Program, University of California, San Francisco; Department of Bioengineering and Therapeutic Sciences, University of California, San Francisco

**Keywords:** deep learning, cell tracking, cell migration, human pluripotent stem cells, morphogenesis

## Abstract

Lineage tracing is a powerful tool traditionally used in developmental biology to interrogate the evolutionary time course of tissue formation, but the dense, three-dimensional nature of tissue limits the ability to assemble individual traces into complete reconstructions of development. Human induced pluripotent stem cells (hiPSCs) enable recapitulation of various aspects of developmental processes, thereby providing an *in vitro* platform to assess the dynamic collective behaviors directing tissue morphogenesis. Here, we trained an ensemble of independent convolutional neural networks to identify individual hiPSCs imaged via time lapse microscopy in order to generate longitudinal measures of individual cell and dense cellular neighborhood properties simultaneously on timescales ranging from minutes to days. Our analysis reveals that while individual cell parameters are not strongly affected by extracellular microenvironmental conditions such as pluripotency maintenance regime or soluble morphogenic cues, regionally specific cell behaviors change in a manner predictive of organization dynamics. By generating complete multicellular reconstructions of hiPSC behavior, our cell tracking pipeline enables fine-grained understanding of developmental organization by elucidating the role of regional behavior stratification in early tissue formation.

## Introduction

In the developing embryo, individual cells undergo a sequence of cell fate transitions and migration events to cooperatively form the tissues and structures of the organism. Cell tracking techniques based upon high resolution imaging have been used to trace cell lineage and describe the emergent patterns of embryogenesis across multiple model organisms,^1–3^ including the early human pre-implantation embryo.^4,5^ However, automated tracking of cell migration within whole embryos *in vivo* has been limited to small organisms such as *C. elegans*^6^ due to the difficulty of identifying and tracking cells in the densely crowded multicellular environment of the developing embryo.

Researchers frequently address the problem of density by employing sparse labeling of cells, either by only tracing cells of a single lineage,^7,8^ or by detecting transcriptional^9^ or morphologic distinctions between cells.^10^ Similarly, when analyzing cell behavior *in vitro,* experimental limitations such as mechanical confinement to one dimensional tracks, ^11^ or sparse labeling^12^ have been required to accurately track individual cells, limiting the ability of these systems to monitor multicellular tissue behaviors with comprehensive single cell resolution.

Self-organizing developmental processes are often initiated by small founder populations within a larger population of physically inter-connected cells, as in the case of classic Turing patterns.^13^ Similar multicellular organizational events have been observed *in vitro* with human induced pluripotent stem cell (hiPSC), revealing their heterogeneous differentiation potential due to global positional cues,^14^ cell population boundaries,^12^ or cell-cell interactions.^15^ In particular, because cell fate and function are strongly impacted by local interactions within multicellular networks,^16–18^ coordinated morphogenic processes exhibit scale-free connectivity (i.e. at multiple scales, cell behavior is coordinated through a central hub of influential cells),^19^ indicating that small populations of cells, by establishing highly connected organizing centers, can exert a large impact on the final composition of the developing tissue.^20,21^ Sparse labeling approaches inherently under-sample these rare populations, highlighting the need for dense cell tracking algorithms to definitively identify the origins and quantify the behaviors of organizers.

Recent advances in machine learning, in particular in deep neural networks, have demonstrated superhuman performance at image segmentation, revolutionizing the field of computer vision.^22,23^ Several classes of convolutional neural nets (CNNs) have been developed specifically to perform dense cell segmentation^24^, based upon different architectures such as autoencoders,^25^ U-nets,^26–28^ or variants of the Inception architecture.^29,30^ Each architecture offers distinct trade-offs between cell segmentation accuracy, training efficiency, noise robustness, and computational complexity, with sub-optimal network choice leading to reduced tracking quality and poor capture of cell behavior. While cell tracking algorithms have historically been assessed through head-to-head competitions^31,32^, the potential advantage of combining complementary techniques for cell localization and tracking has been rarely employed.

In this study, we overcame the challenge of dense cell tracking by developing an ensemble of three neural networks (FCRN-B,^24^ Count-ception,^30^ and a Residual U-net^27^) to localize each individual cell nucleus in an hiPSC colony. Nuclei displacements were then connected between sequential frames of a time series, enabling high spatiotemporal resolution of hiPSC behaviors over relevant developmental time scales of hours to days. This dense cell tracking pipeline revealed distinctive cell behaviors based on location within the colony, cell heterogeneity, and response to extracellular signaling molecules. Long-term cell tracking in combination with immunostaining for lineage markers, enabled tracking of the differentiation history of colonies with singlecell resolution. The whole-colony tracking and analysis pipeline revealed radially stratified shifts in cell migration speed and cell packing density in hiPSC colonies in reaction to changes in culture conditions. Changes in cellular behavior were detected at the local cell neighborhood level in response to differentiation induced by externally applied morphogens, enabling real-time identification of local organizing centers (~10-20 cells) that precede tissue-scale morphogenic events. By detecting rare organizational events, our computational cell tracking pipeline allows for a more comprehensive dynamic understanding of the multicellular principles of morphogenesis, which can empower more refined control of organoid and engineered tissue development.

## Results

### Manual annotation of cell migration in hiPSC colonies

Human iPSCs form dense, multilayered colonies *in vitro* with indistinctive boundaries between cells when using common phase imaging, pan-cytoplasmic, or pan-nuclear staining techniques. To establish a baseline for cell localization quality, a series of heterogeneously labeled colonies were generated by mixing wild type hiPSCs with an hiPSC-derived cell line expressing a nuclear GFP fluorescent label (Lamin-B::GFP) at ratios of 9:1 (10% labeled), 7:3 (30% labeled) or 0:10 (100% labeled). While maintaining the cells in pluripotency media, mixtures were force aggregated in microwells,^15^ allowed to reattach to tissue culture plates, and then imaged every five minutes for six hours to generate a set of frames for annotation (Figure 1A, B). Seven individual human annotators selected the center of every GFP positive cell nucleus in 12 sequential pairs of 500-cell colonies containing 10%, 30%, or 100% Lamin-B::GFP iPSCs presented in randomized order. A spatial average of all seven annotation sets was calculated using k-means clustering to generate a ground truth human consensus segmentation for each frame (Figure 1B).

**Figure 1.**
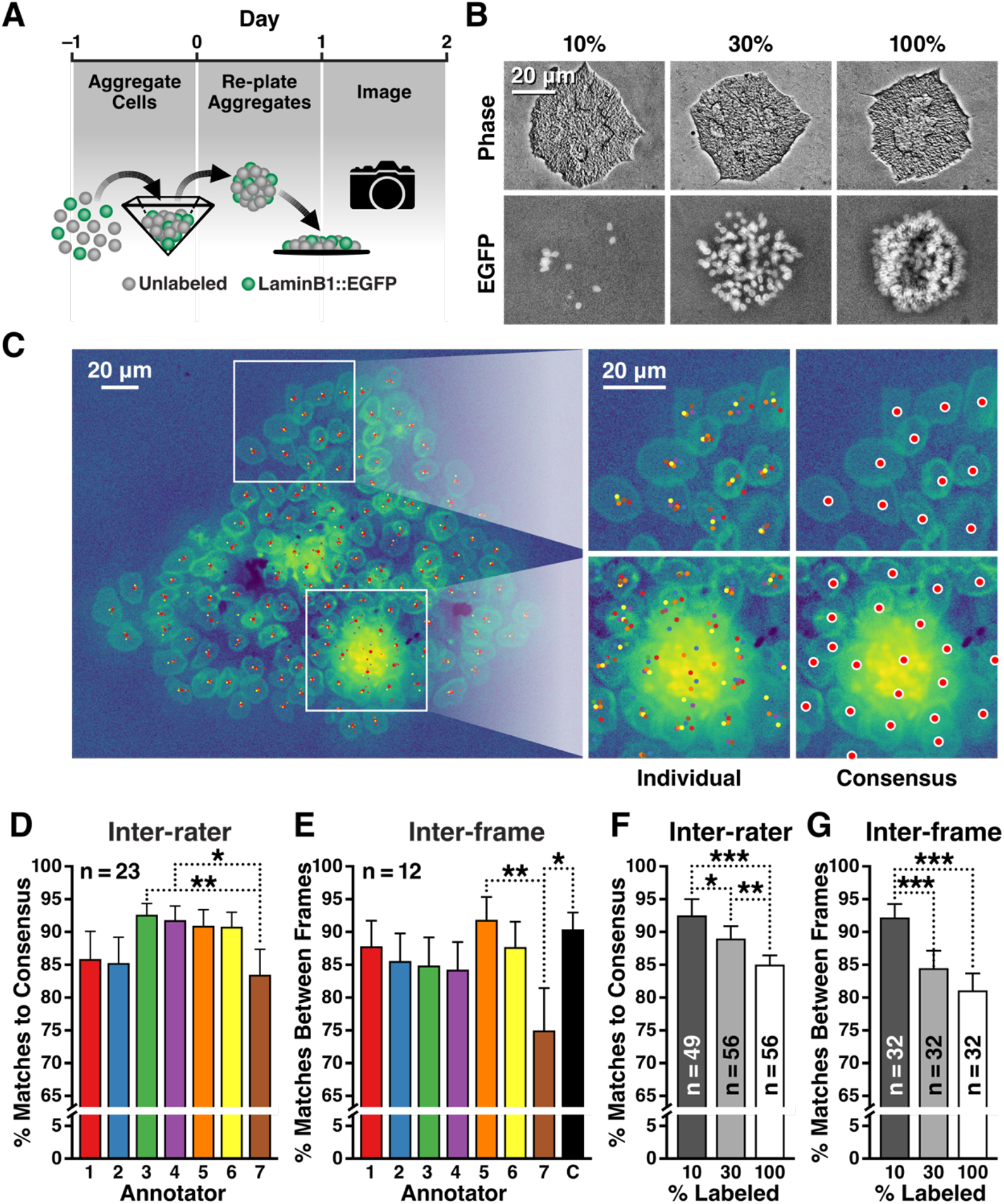
Quality of Manual Tracking Plateaus with Increasing Density of Labeled Cells. A. WTC11 and LaminB1::GFP cells were seeded into microwells, generating mixed aggregates with defined ratios of each population, then the aggregates were re-plated to form colonies. B. Annotators selected all cells in the colony, with high consistency in sparse regions, but lower agreement in dense regions. C. Individual annotator accuracy for each image was compared to the consensus for all images (n=23) with only annotator 7 different from any other rater (* p < 0.05, ** p < 0.01). D. Accuracy segmenting the same cell across sequential frames was assessed for all image pairs (n=12) but only annotator 7 was less repeatable than other annotators (* p < 0.05, ** p < 0.01). E. Colonies with 10%, 30%, and 100% LaminB1::GFP labeled cells were formed with labeled cells dispersed throughout the colony. F. Annotators were less accurate as compared to consensus on 100% labeled colonies than on 10% and 30% colonies (** p < 0.01, *** p < 0.001). G. Segmenting the same cell across sequential frames was also less repeatable in 100% labeled colonies vs 10%

Human annotators were scored using a ratio of selected nuclei within a 5 *μm* radius from the consensus cell, divided by the total number of expected cells, missing cells, and incorrectly selected cells (true positives divided by all positives plus any false positives). The average individual rater reliability (IRR) was 88.5% (± 7.9 %) with a minimum of 83% and a maximum of 93% (Figure 1C). As a second comparison, the individuals and human consensus were rated on their ability to select the same cell twice in pairs of sequential frames. Average inter-frame reliability (IFR) was 85.8% (± 7.7%) with a minimum of 75% and a maximum of 92% (consensus 90.2% ± 4.8%) (Figure 1D). To assess how increasing label density impacted cell detection, performance metrics were stratified according to colony labeling density (Figure 1E). As expected, human annotators exhibited maximal IRR and IFR when evaluating colonies with the lowest percentage of GFP+ cells (i.e. 10%), with performance significantly declining for colonies containing higher proportions of GFP+ cells (30% and 100%).

### Ensemble deep neural network segmentation of dense hiPSC colonies

To determine how deep neural networks compare to human segmentation performance, a diverse array of independent cell segmentation network architectures was selected from recent literature (Figure 2Ai) and compared to the human annotator baseline as well as to the prediction of an ensemble of the selected architectures (Figure 2Aii). Five different neural net architectures were compared, including two networks with VGG-like architecture (FCRN-A and FCRN-B^24^), two U-net architectures (U-net^28^ and Residual U-net^27^), and an Inception-inspired network (Count-ception^30^). Each neural network was trained to segment the GFP images of 10%, 30%, and 100% labeled colonies by predicting a cone-shaped probability around the human annotated center of each nucleus. Despite architectural differences, all neural networks exhibited comparable average performance, segmenting the data with a receiver operating characteristic (ROC) area under the curve (AUC) of 0.86 or better (Figure 2B). Although no individual neural network was able to equal human segmentation of 100% GFP-labeled colonies, an ensemble of the three highest performing networks surpassed human cell localization of fully-labeled colonies (Figure 2Aii, B-D). The primary variation between neural networks was due to spatial performance differences at the center or edge of individual colonies (Figure 2E).

**Figure 2.**
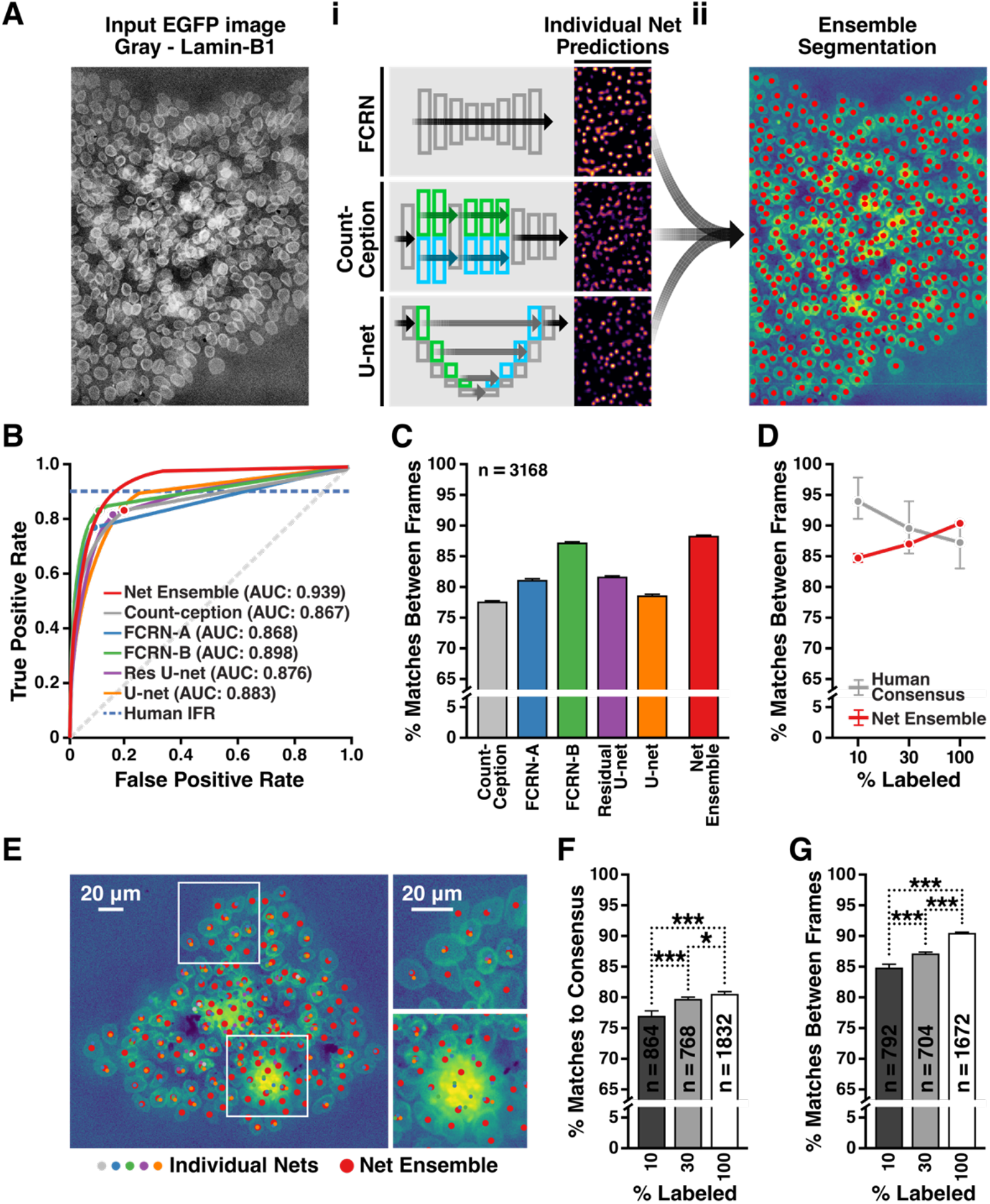
Heterotypic Neural Net Ensembles Generate Human Quality Segmentations. A.i. Individual images were segmented by one of several neural net architectures. producing probability maps localizing the center of each LaminB1::GFP labeled nuclei. ii. A weighted average of these maps from three different architectures (FCRN-B, Residual U-net, Count-ception) was used to produce the consensus segmentation. B. The true positive and false positive rate was calculated for each segmentation over the range of probability map thresholds between 0 and 1 and then the area under the curve (AUC) calculated for each architecture. C. Repeatability of cell detections between frames was calculated for the entire training set (n=3,168, p < 0.001 vs all single neural nets). D. Repeatability of cell detections was stratified by percent labeling and compared to the human annotator consensus, (* p < 0.05, 30% and 100% not significant). E. Representative image depicting individual and net ensemble detection ability where different colored dots indicate the peak probability of a cell as predicted by each neural net architecture. F. The agreement between net ensemble predicted labels and the human annotated data set was assessed for each label percentage (* p < 0.05, *** p < 0.001) G. The repeatability of the net ensemble detections over time was also compared across label percentages (*** p < 0.001).

Compared to the human annotator baseline, cell segmentation performance varied greatly between networks, with the two U-net architectures agreeing least with human annotators, whereas FCRN-B and the ensemble agreed most often (Figure S1). However, ROC AUC (Figure 2B) and effect size were often indistinguishable between similar architectures such as FCRN-A and FCRN-B (Cohen’s *d* = 0.11) or U-net and residual U-net (*d* = 0.02). Segmentation speed varied widely between architectures (from 23 to 288 milliseconds per frame), but because the ensemble network was composed of several of the faster architectures, generating the composite segmentation was only 16.0% slower than using U-net only (± 2.5% slowdown; Figure S2). In contrast to human annotators, neural net IRR and IFR segmentation accuracy improved with increasing label density (3.6%±0.4% and 5.7%±0.4%, Figure 2F,G respectively).

### Individual cell tracking of pluripotent stem cell behavior

Individual frame segmentations were initially combined using a nearest neighbor linkage between frames to create cell tracks covering the center, middle, and edge regions of each colony (Figure 3A, blue, gold, and red regions respectively), enabling construction of whole colony traces for all cells in 100 % GFP+ colonies over the entire time series (Figure 3B). However, segmentation uncertainty at the individual cell level (e.g. a 95% accurate classifier will fail to detect a cell approximately once every 20th frame) led to artificially shortened tracks separated by short gaps of 1-5 frames. To reduce random breakages, a second linking step was added to combine tails of track fragments across gaps of up to 5 frames, using the motion of the local cellular neighborhood to interpolate any missing cell positions. Neighborhood interpolation significantly increased track fragment lengths (from average coverage of 21.5% of the time series length to 33.5%, Figure S3), bringing track fragment counts closer to the expected cell count based on cell seeding number and extrapolated growth rate - from 482 to 836 individual cells, with 1,000 expected (Figure S4).

**Figure 3.**
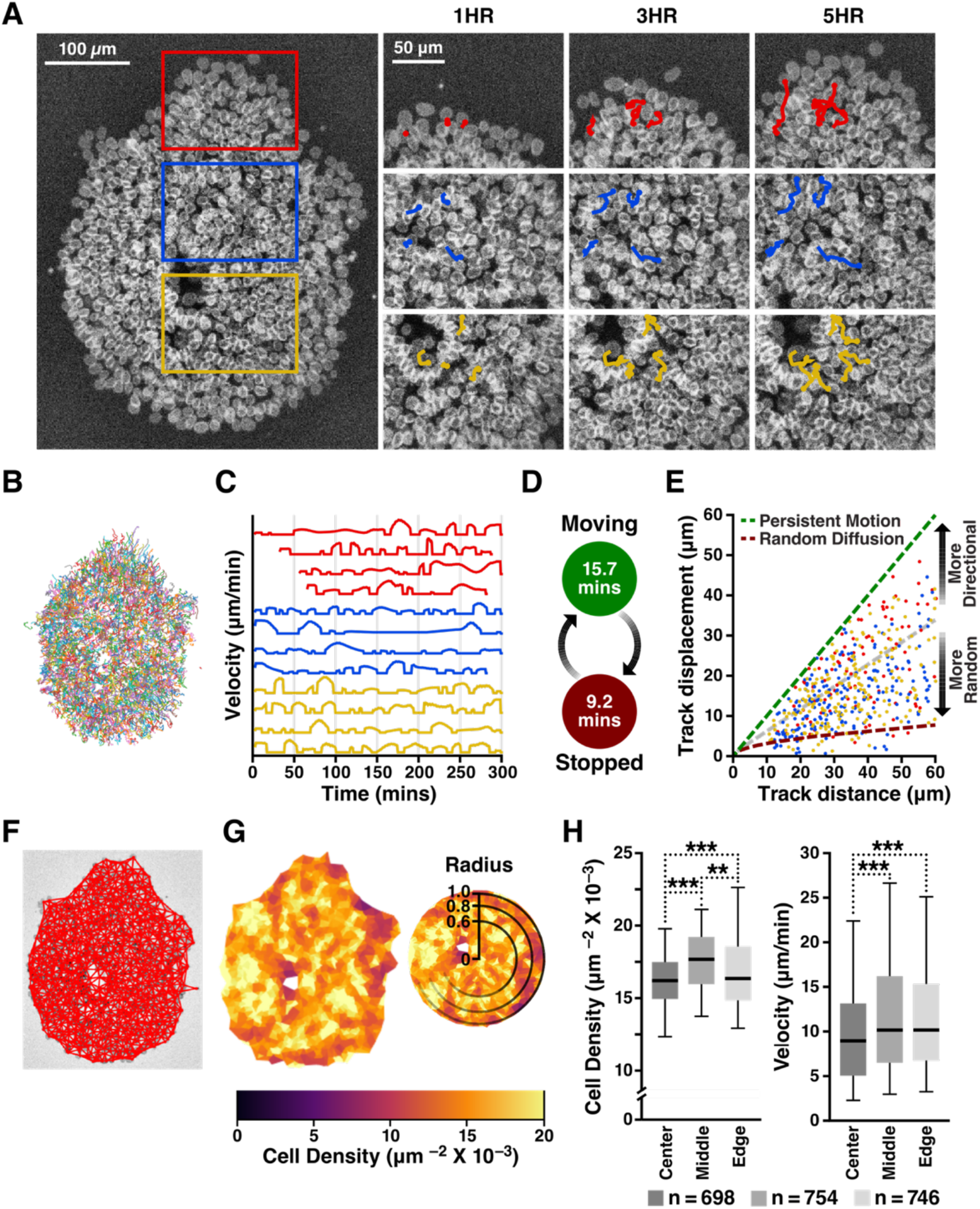
Spatio-temporal linkage of detections enables long term single cell tracking. A. Individual detections were linked across frames forming long tracks that spanned the entire time series. Depiction of example colony where regions were identified as center, middle, edge (colors). B. Dense track map created by linking detections covering the entire time series. C. Trace plot of example cell velocity tracks across colony locations (colors) D. Proposed two state model of alternating active migration and quiescence fit from average active and stopped periods. E. Distribution of the ratio of total track displacement to total track distance where colored dots represent individual tracks from the center, middle, and edge regions and dotted lines show the theoretical curves for persistent migration (dark green) and random diffusion (dark red) F. Delaunay triangulation depicted across an example colony to calculate cell neighborhoods, G. Average inverse area of Delaunay triangles around each cell (cell density) depicted on the example colony, and projected onto the unit circle. H. Quantification of cell density and velocity across the colony region identified by projecting triangulated cell position onto rings of the unit circle (** p < 0.01, *** p < 0.001).

To understand how individual cell behavior contributes to colony spreading and density, we calculated persistence of cell migration by locating regions of each track where the direction of cell motion changed by less than 5 degrees per minute. Most cell tracks displayed clear binary switching between persistent migratory and stationary behavior (Figure 3C), with a mean active period of 15.7 minutes (±12.7 minutes) followed by a quiescent period of 9.2 minutes (±6.8 minutes), similar to the cyclic migration behavior observed in *E. coli*^33^ and eukaryotic cells^34^ that can be attributed to the interaction between local polarizing cues and global inhibition of directional migration (Figure 3D). The active migration period was highest at the edge of colonies, and lowest at the center, while the quiescent period did not differ between colony regions (Figure S3-3).

To measure the extent to which individual cells traveled directionally or diffused randomly, we calculated the ratio of track displacement-to-distance, where a value of 1.0 represents travel in a straight line, lower values an increasingly curved trajectory, and 0.0 a path that ultimately returns to its origin. Although cell tracks covered a broad range between purely directional and random diffusion, there was no difference in directionality of motion between cells at the center, middle or periphery of the colony (Figure 3E, blue, gold and red points respectively). Finally, to identify coordinated movement between neighboring cells, we calculated correlation between each cell’s velocity profile and its immediate neighbors. In the center of colonies, nearest neighbors had uncorrelated velocity profiles (Pearson’s R = 0.00298, std 0.0757), whereas cells near the periphery demonstrated much higher correlation (R = 0.118, std 0.383), suggesting that observed peripheral spreading results from multi-cellular collective migration, as has been shown previously in models of collective migration.^35,36^

To analyze the dynamic behavior of iPSC colonies, a graph structure of each colony was created using Delaunay triangulation (Figure 3F). Based on the triangulation, individual cell area was estimated using the average of all triangles surrounding a cell within a maximum link distance threshold of 50 *μm* (Figure 3G). The entire colony mesh and all cell measurements, such as density or velocity, were mapped onto the unit circle, then separated into three rings of equal area corresponding to the center, middle, and periphery of the colony (Figure 3G). In pluripotent colonies, cells in the center region were packed more densely relative to the middle and peripheral bins (p = 1.82*10^−10^ and 1.56*10^−11^, respectively) (Figure 3H), suggesting local crowding effects contribute to radial inhomogeneities in cell packing in hiPSC colonies. In contrast, cells in the middle and peripheral bins moved faster than cells in the center (p = 2.28*10^−5^ and 2.93^−4^ respectively, Figure 3H), demonstrating an edge-biased cell migratory phenotype and suggesting that colony compaction may play a role in hiPSC colony spreading, as has been reported for migration of other epithelial cells.^35,36^

### Packing and migratory behaviors of undifferentiated pluripotent stem cells

To interrogate the heterogeneous behavior of hiPSC colonies, we compared standard pluripotency maintenance conditions using the CNN tracking algorithm. First, we compared the effect of colony size on single cell behavior by forming colonies of either 100 or 500 cells (Figure 4Ai). The average cell density and travel distance of 100-cell colonies were more similar to those of the edge of 500-cell colonies than to the center, suggesting that 100-cell colonies uniformly exhibit a similar phenotype to the edge of 500-cell colonies (Figure 4Aii). At both colony sizes, cells at the edge displayed higher travel distance and migration speeds than those at the center (Figure 4Aiii,iv). 100-cell colonies were more uniform in both density and cell distance traveled, with both measures closer to the cell density and travel values for the edge of 500-cell colonies. The transition in phenotype from edge-like to center-like cells as confluency increases may account for the observed sensitivity of hiPSC pluripotency and differentiation to cell plating density^37^ and colony size.^14^

**Figure 4.**
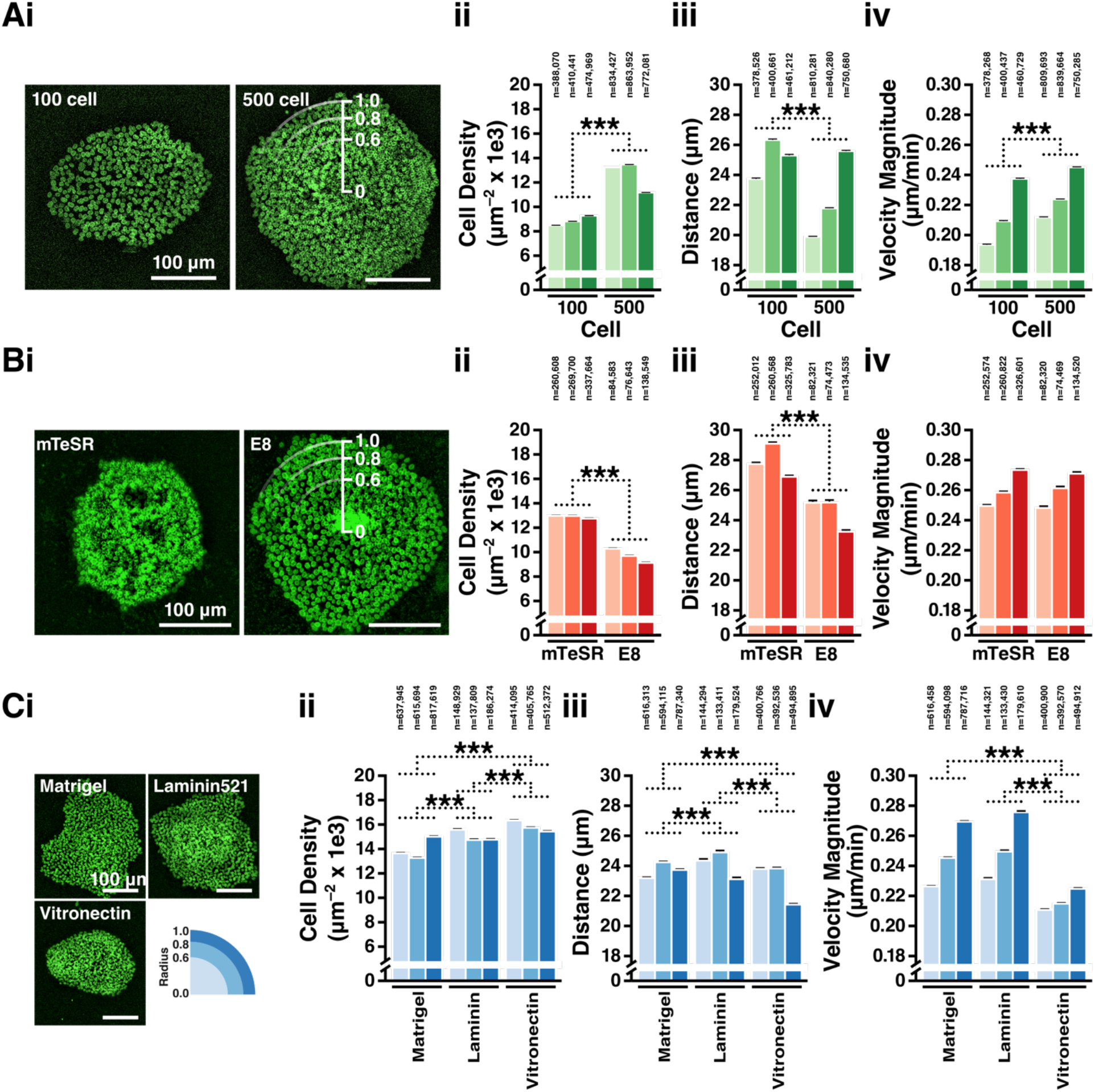
Basal culture conditions change cell packing density and migratory behavior. A.i. Example images of colonies with 100 or 500 starting cells. Comparison of 100 cell and 500 cell colonies stratified by colony region for: ii. average cell density (*** p < 0.001), iii. average total cell distance traveled in 6 hours (*** p < 0.001) and iv. average instantaneous cell velocity (*** p < 0.001). B.i. Example images of colonies generated from cells cultured in mTeSR or E8. Comparison of mTeSR and E8 colonies stratified by colony region for: ii. average cell density (*** p < 0.001), iii. average total cell distance traveled in 6 hours (*** p < 0.001) and iv. average instantaneous cell velocity (not significantly different). C.i. Example images of colonies adhered to either Matrigel, Vitronectin, or rLaminin521 coated plates. Comparison of Mtrigel, Virtonectin and rLaminin521 colonies stratified by colony region for: ii. average cell density (*** p < 0.001), iii. average total cell distance traveled in 6 hours (*** p < 0.001) and iv. average instantaneous cell velocity (*** p < 0.001, Matrigel and rLaminin521 not significantly different)

Next, we explored the effect of pluripotency maintenance media on colony behavior by comparing the effect of passaging hiPSCs in mTeSR or E8 media (Figure 4Bi). Colonies cultured in mTeSR were more compact with frequent formation of multi-layered structures and low-density regions in the center of the colony, while colonies cultured in E8 were uniformly flat with lower cell packing density (Figure 4Bii). Individual cells within colonies cultured in E8 traveled less overall (Figure 4Biii). Despite structural differences, cell migration velocities between the two conditions only differed slightly (p = 0.012, d = 9.44*10^−3^), indicating that the density shift could not be solely attributed to differences in cell motility between the two conditions (Figure 4Biv).

Finally, we interrogated changes in colony phenotype due to commonly used adhesive extracellular matrices, which have been shown to have a cell-ECM strain-mediated effect on hiPSC morphology, behavior, and differentiation potential.^38^ hiPSC aggregates were allowed to adhere onto either Matrigel, Vitronectin, or recombinant Laminin 521 (rLaminin, Figure 4Ci). Cell adhesion was much lower on rLaminin, with only 47.2% of aggregates adhered after 24 hours vs 91.7% on Matrigel and 97.2% on Vitronectin. Cells in adherent colonies on both rLaminin and Vitronectin had higher cell density than on Matrigel (Figure 4Cii), while cells on Matrigel and rLaminin spread more than on Vitronectin (Figure 4Ciii). Cells on Vitronectin had lower migration velocities, and much lower difference between center and edge migration velocities than either Matrigel or rLaminin (Figure 4Civ). hiPSC behavior on Matrigel and rLaminin were very similar for both cell migration distance and migration velocity, however stratifying the colonies by radius revealed that colonies plated on Matrigel were 11.1% less dense in the center. hiPSCs at the periphery of colonies grown on Vitronectin traveled only 92.7% of the distance for those on the edge of Matrigel or rLaminin colonies, and cells in Vitronectin colonies uniformly moved more slowly than those on other matrices, leading to more compact colony morphology overall. These results suggest that changes to substrate can subtly alter the local strain environment within a pluripotent stem cell colony, providing a mechanism to modulate peripheral migration and cell packing within hiPSC colonies.

Through dynamic characterization of hiPSC behavior, our tracking pipeline revealed that hiPSCs display a wide variety of heterogeneous behaviors while maintaining pluripotency. In particular, cells at the periphery of colonies exhibit a distinct phenotype from those in the center. Media environment and substrate can modulate both static and dynamic aspects of the edge and center phenotype. However, static snapshots of colony configuration, such as cell density, do not predict dynamic cell behaviors such as cell migration distance or velocity. Since both static and dynamic cell behaviors prime hiPSCs towards particular differentiation trajectories,^14,38–40^ dynamic assessment of whole colony behavior is necessary to illuminate the scope of hiPSC heterogeneity in pluripotency and predict priming during differentiation.

### Lineage tracing of cell fate decisions during early morphogenic induction

We next assessed changes in hiPSC behavior during early lineage specification by employing our tracking pipeline to analyze differentiation protocols used to induce combinations of all three germ layers. Previous work has shown that multi-cellular annular ring patterns form during tri-lineage differentiation,^14^ but the dynamic changes to cell migration behavior during ring formation have not been described. In addition, protocols to induce either mesendoderm^37^ or neuroectoderm^41^ have been reported, but whether those direct differentiation protocols induce similar dynamic transformations to those that occur during tri-lineage differentiation is not known. To monitor the transition from pluripotent cells to differentiating germ layers, a critical 24-hour morphogenic window was identified for each differentiation protocol for further exploration.

In the BMP4-induced trilineage protocol (Figure 5Ai), colonies adopted a round morphology 24 hours after BMP4 treatment with relatively uniform velocity and cell density, consistent with undifferentiated colonies (Figure S5). Approximately 32 hours after induction, cells across the colony slowed in migration velocity, except for a ring of cells at ~50% of the colony radius which maintained similar velocity to undifferentiated cells (Figure S6A). In the center of the colony, cell density was constant for the entire period of observation, however, the periphery of the colony also began to rapidly decrease in cell density about 32 hours post-induction, with a dense plateau of cells forming at approximately 50% of colony radius, consistent with previous reports^14,42^(Figure 5Aiii, Figure S7A). All three germ lineages formed by 48 hours, with OCT4+ cells in the center ring (SOX2-, EOMES-, presumptive endoderm), EOMES+ cells in the middle (presumptive mesoderm), SOX2+ cells at the colony edge (OCT4-, EOMES-, presumptive ectoderm), and the periphery of the colony negative for all three markers (Figure 5Aiii). The peak of EOMES expression corresponded with both the maximum of cell migration velocity and the transition from high to low cell density, suggesting that the mesoderm ring acts as a migratory barrier between ectoderm and endoderm, enabling the physical phase separation of the colony into three distinct germ layers, analogous to gastrulation^14,21,42^.

**Figure 5.**
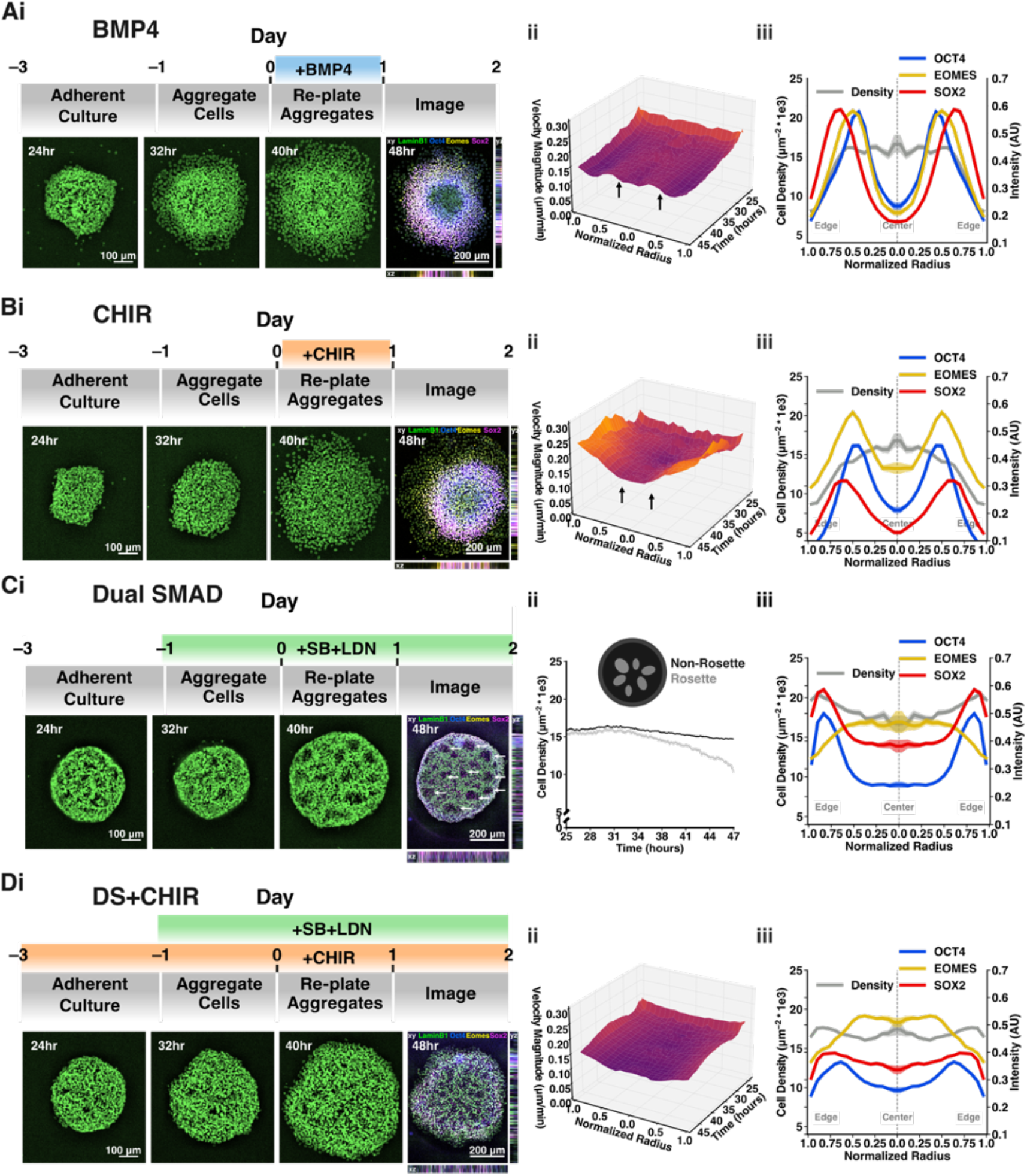
Whole Colony Analysis Reveals a Density Signature of Multi-lineage Differentiation. A.i. Treatment timeline and example time course of colony treated with BMP4 with example images at 24, 32, and 40 hours post re-seeding (HPR) and fixed and stained image of the same colony at 48 HPR. ii. Surface plot of temporal evolution of average instantaneous cell velocity over BMP4-treated colonies projected on to the unit circle (n=16 colonies). iii. OCT4, SOX2, and EOMES expression profiles and average cell density profile at 48 HPR projected onto the unit circle in BMP4-treated colonies (n=16 colonies). B.i. Treatment timeline and example time course of colony treated with CHIR with example images at 24, 32, and 40 HPR and fixed and stained image of the same colony at 48 HPR. ii. Surface plot of temporal evolution of average instantaneous cell velocity over CHIR-treated colonies projected on to the unit circle (n=16 colonies). iii. OCT4, SOX2, and EOMES expression profiles and average cell density profile at 48 HPR projected onto the unit circle in CHIR-treated colonies (n=16 colonies). C.i. Treatment timeline and example time course of colony treated with Dual SMAD inhibition at 24, 32, and 40 HPR and fixed and stained image of the same colony at 48 HPR with rosettes highlighted (white arrows). ii. Temporal evolution of average cell density inside and outside of rosettes (n=16 colonies). iii. OCT4, SOX2, and EOMES expression profiles and average cell density profile at 48 HPR projected onto the unit circle in Dual SMAD inhibition-treated colonies (n=16 colonies). D.i. Treatment timeline and example time course of colonies treated with both Dual-SMAD inhibition and CHIR pre-treatment, at 24, 32, and 40 HPR with fixed and stained image of the same colony at 48 HPR. ii. Surface plot of temporal evolution of average instantaneous cell velocity over DualSmad+CHIR-treated colonies projected on to the unit circle (n=16 colonies). iii. OCT4, SOX2, and EOMES expression profiles and average cell density profile at 48 HPR projected onto the unit circle in Dual SMAD+CHIR-treated colonies (n=16 colonies).

Treatment with the WNT activator CHIR is commonly used to induce differentiation of mesoderm.^37^ Tall, multilayered colonies (average 61.4 ± 10.7*μm*) formed after 24 hours of 12 μM CHIR treatment, but by 48 hours a secondary flat epithelial ring expanded radially out from the colonies, ultimately forming a stratified colony similar to that induced by BMP4 (Figure 5Bi). Unlike in BMP4-treated colonies, CHIR-treated cells at the colony periphery increased in migration speed by 50%, with individual cells at the periphery of the colony undergoing EMT and traveling beyond the field of view (Figure S6B). Similar to BMP4 treatment, the central region maintained cell density similar to untreated colonies, while the middle and outer compartments rapidly decreased in density (Figure 5Biii, S7B). OCT4, SOX2, and EOMES were detected in all colonies, but levels of OCT4 and especially SOX2 were lower with CHIR than in BMP4 treated colonies, consistent with early CHIR induction directing differentiation towards mesoderm and away from neuroectoderm (Figure 5Biii). Again, the peak of EOMES expression occurred at ~50% of the colony radius and corresponded spatially to the transitions between low to high velocity and high to low density, respectively. The direct comparison of CHIR and BMP4 induced differentiations demonstrates that limited numbers of similar static snapshots of colony structure can mask distinctive cell behaviors that can indicate divergent differentiation trajectories of pluripotent cells.

Neuro-ectoderm directed colonies remained behaviorally indistinguishable from untreated colonies through the first 48 hours of dual SMAD inhibition. However, starting at 60 hours after treatment, small rosettes of approximately 20 cells began to form ring structures that expanded continuously for the remaining 12 hours of imaging (Figure 5Ci). Between 6 and 18 rosettes formed per colony (mean 10.1 ± 2.3) with a mean rosette diameter of 64.2 ± 21.1*μm*. Rings consisting of regions of lower cell density began to appear 36 hours after plating, with ring diameter expanding at a rate of 2.58 ± 0.51*μm/hour*, and a mean center-to-center spacing between rings of 124.0 ± 27.5*μm* (Figure 5Cii). Average cell density was slightly higher at the periphery of colonies, corresponding to higher expression of both OCT4 and SOX2 (EOMES-, potentially undifferentiated cells), while the center of the colonies expressed high SOX2 and low OCT4 (presumptive neuroectoderm, Figure 5Ciii). EOMES expression was slightly elevated in the center of the colonies, but overall EOMES was rarely detected compared to BMP4 or CHIR differentiation. None of the three lineage markers appeared to be specifically localized to the ring structures. Addition of CHIR pre-treatment^43^ to the dual SMAD neuro-ectoderm protocol completely abrogated the formation of rosettes (Figure 5Di). CHIR pre-treated neuro-ectoderm colonies were indistinguishable from untreated colonies in both their uniform velocities and radial distribution of cell densities (Figure 5Dii and S6D, S7D, respectively). CHIR treatment elevated expression of EOMES, and suppressed expression of both SOX2 and OCT4, likely delaying the commitment of cells to neuroectoderm fates, consistent with its previously reported activity.^43^ By monitoring the trajectories of differentiating colonies at single cell resolution, morphogenic signatures were detected at both the local cell neighborhood and colony-wide levels, thereby enabling quantitative measurement of the comprehensive dynamics of multicellular organization and subtle yet distinctive differences in cell behavior that distinguish between independent differentiation protocols.

## Discussion

Single cell analyses have highlighted the intrinsic heterogeneity present in virtually all multi-cellular populations. Complementary approaches, such as automated cell lineage tracing and single-cell RNA sequencing, have enabled fine-grained spatio-temporal quantification of diverse and robust developmental processes^6,7,44^. Understanding the dynamic behavior(s) of pluripotent stem cells in response to environmental factors can similarly clarify the effects of multicellular structure and environmental factors on the behavior and ultimate fate of individual cells within developing tissues and organs *ex vivo*. To assess how organization arises from the collective action of individual cells, we developed a dense cell tracking platform to analyze time lapse imaging of hiPSC colonies with high spatiotemporal precision. Using the resulting quantitative measures of cell behaviors, we identified signatures of multicellular organization at the single cell, local neighborhood, and whole colony scale, demonstrating that hiPSC behaviors are influenced by short distance interactions between neighboring cells that propagate into global effects throughout an entire colony of 100’s of cells and more. While many of the measured cell-intrinsic properties were relatively constant under pluripotent culture and early differentiation conditions, we found that the local cell neighborhood responds in characteristic ways to different external stimuli. Changes in cell-cell interactions are orthogonal to stem cell pluripotency,^12,38^ but can impact the sensitivity of hiPSCs to morphogenic cues,^39^ and thus may be a critical determinant in pre-patterning of cells to different cell fate decisions. The ability to specifically modulate cell-cell interactions through modification of culture conditions or genetic engineering provides new strategies to pre-pattern and control colony structure and subsequent differentiation trajectories.^39^ Furthermore, our live cell monitoring platform during early differentiation allows for non-destructive assessment of regional changes in cell fate, providing a critical first step towards feedback-control of hiPSC differentiation.

In this paper, we applied our tracking system to resolve human pluripotent morphogenesis evolution at single-cell resolution to the maintenance of hiPSCs and early differentiation, but it can be used more generally to quantify multicellular structure with either static or time-lapse microscopy of any cell line. Quantitative comprehensive characterization of cellular neighborhood dynamics will provide a robust approach to interrogate the effects of multicellular interactions among a broad range of cell types across many species, and will provide novel metrics to assess the fidelity of stem cell models to recapitulate developmental processes in a tissue context *ex vivo.* Ultimately, extracting unbiased cell dynamics from *in vitro* time-lapse imaging enables new insights into the complex processes underlying multicellular organization and morphogenesis.

## Methods

### hiPSC culture

The hiPSC cell lines Wild-Type C11 was provided by the Conklin Lab and the Allen institute Lamin-B1 EGFP line (AICS-0013) was provided by Coriel. Both lines were cultured in feeder-free media on growth factor reduced Matrigel (BD Biosciences) and fed daily with mTeSR^TM^-1 medium (STEMCELL Technologies)^45^. Stem cells were dissociated to single cells using Accutase (STEMCELL Technologies) and passaged at a seeding density of 12,000 cells per cm^2^. Rho-associated coiled-coil kinase 1 (ROCK-1) inhibitor, Y-276932 (10 *μ*M; Selleckchem) was added to the media for the first 24 hours after passaging to promote survival^46^. Starting with a base cell line cultured in mTeSR, cells were migrated to E8 (Gibco) by first culturing for one passage in a 1:1 mixture of E8 and mTeSR, followed by a minimum of two passages in E8 before evaluation.

### Force aggregation of colonies

Cell aggregates consisting of 250 cells were generated using 400×400 *μ*m PDMS microwell inserts in 24-well plates (975 microwells per well).^15^ Dissociated cultures were resuspended in their respective maintenance media supplemented with Y-276932, mixed at the required cell ratio and concentration (250 cells/well), added to the microwells, and centrifuged (200 RCF). After 24 hours of formation, aggregates were transferred to ibidi slides (12 uWell format) or optically clear 96-well plates (Corning), coated with a substrate, and seeded at 10 aggregates/well or 18 aggregates/*cm*^2^ and allowed to spread into colonies.

### Substrate coating protocol

To promote aggregate attachment, ibidi slides were coated with growth factor-reduced Matrigel (80 *μg/mL,* BD Biosciences), vitronectin (10 *μg/mL,* Sigma Aldrich) or recombinant human laminin 521 (*10μg/mL* rLaminin, Corning). Wells were uniformly coated using 125 *μ*L/well (223 *μL/cm*^2^) and incubated at 37 C following manufacturer’s recommendations. Matrigel was incubated for 16 hours, while vitronectin and rLaminin were both incubated for 1 hour. Following manufacturer recommendation, rLaminin wells were additionally washed three times using cell culture grade water.

### Differentiation protocol

Recombinant BMP4 (R and D Systems) was added to mTeSR at 50 ng/mL for 24 hours, starting 24 hours after aggregate seeding to induce a trilineage differentiation,^14^ followed by 24 hours of imaging in mTeSR alone. CHIR differentiation was performed by adding 12 *μM* CHIR-99021 (Selleck Chemicals) to mTeSR for 24 hours, starting 24 hours after seeding, followed by 24 hours of imaging in mTeSR only. Dual SMAD inhibition was performed by adding both 10 *μM* SB-431542 (GlaxoSmithKline) and 0.2 *μ*M LDN-193189 (Stemgent) to mTeSR^41^ during force aggregation, and maintained for 72 hours through colony adhesion and imaging. CHIR pre-treatment during dual SMAD inhibition was achieved by adding 2 *μM* CHIR-99021 to mTeSR starting 48 hours before force aggregation, and continued through imaging (5 total days of treatment)^43^.

### Time-lapse imaging

Initial mixing studies were performed on an incubated inverted Axio Observer Z1 (Zeiss) microscope using an AxioCam MRm (Zeiss) digital CMOS camera at 20x magnification (NA 0.8, 0.323 *μm* x 0.323 *μm* per pixel). Colony positions were mapped using ZenPro software and approximately 30 colonies were imaged each experiment. Colonies were imaged over the course of 6 hours with images taken every 5 minutes.

All subsequent studies were performed on an incubated spinning disk confocal Observer Z1 (Zeiss) using a motorized filter wheel (Yokogawa) and imaged using a Prime 95B (Photometrics) digital CMOS camera at 10x magnification (NA 0.45, 0.91 *μm* x 0.91 *μm* per pixel). Pluripotent colony studies were imaged over 6 hours with images taken every 3 minutes. Differentiation studies were imaged over 24 hours with images taken every 5 minutes.

### Immunofluorescence staining

Within 15 minutes of the conclusion of imaging, slides and plates were washed once with PBS (125 μL/well, 220 *μ*L/*cm*^2^), then fixed for 30 minutes with 4% paraformaldehyde (100 *μ*L/well, 178 *μ*L/*cm*^2^). Cells were permeabilized for 1 hour in 200 *μ*L/well (357 *μ*L/*cm*^2^) of a blocking solution of 5% normal donkey serum, 0.3 % Triton X-100 in PBS. Cells were incubated with the primary antibody for 1 hour in 100 *μ*L/well (178 *μ*L/*cm*^2^) of a solution of 1% bovine serum albumin, 0.3 % Triton X-100 in PBS. Cells were then incubated with the secondary antibody for 1 hour in 100 *μ*L/well (178 *μ*L/*cm*^2^) of a solution of 1% bovine serum albumin, 0.3 % Triton X-100 in PBS.

### Human labeling of data set

To establish baseline human performance on labeling colonies, one annotator labeled the first 12 frames of the time series for 8 colonies each of 9:1 (10% labeled), 7:3 (30% labeled) or 0:10 (100% labeled) wild type:GFP+ mixed colonies (336 frames total). A power analysis was performed, indicating that 8 samples per condition in a 3-way balanced design was required to distinguish between annotator performance when labeling different colony mixture ratios (p <= 0.05 with 80 % confidence of rejecting the null hypothesis).

To produce a validated human data set, 12 pairs of random sequential frames from the original labeled data set were shuffled to obscure the order of the images (for a total of 24 images, 8 per condition). With 50 % probability, each image was horizontally mirrored (13 mirrored, 11 not mirrored), then with 25 % probability, each image was randomly rotated in increments of 90 degrees (6 unrotated, 6 rotated 90 degrees, 9 rotated 180 degrees, 3 rotated 270 degrees). This data set was presented to seven independent annotators using custom software written in Python.

A consensus segmentation was generated using k-means clustering of all annotations on each frame with k equal to the largest number of points selected by any individual annotator. Annotations were added to the consensus if they had at least 3 members in a cluster from unique annotators. This consensus segmentation was used as ground truth to calculate inter-rater reliability (IRR) for each annotator for each frame.

Each frame to frame segmentation accuracy was measured using each annotator’s inter-frame reliability (IFR) on the 12 pairs of images after inverting their transformation. IFR was compared between all pairwise two-sided t-tests with the Bonferroni Holm correction for multiple comparisons.

To evaluate the ability for this segmentation architecture to transfer learning to a different microscope, a second data set of confocal images was segmented by a single human annotator. The first two frames of each of 12 10% labeled colonies, 12 30% labeled colonies, and 8 100 % labeled colonies were segmented in sequence with no crops, flips, or rotations. Inter-frame reliability on this data set was not significantly different from the previous annotations, so this data set was used as a baseline for performance for transfer learning.

### Initial neural net training

Each neural network architecture was implemented using Keras with the Tensorflow back end, with the input field of view for each architecture enlarged to 256×256 to enable fully convolutional segmentation of large images. Neural networks were trained using two NVidea GeForce GTX 1080 GPUs.

A training data set was generated from the initial segmentation by first splitting the images into 80% training, 10% test, and 10% validation folds. Then each fold was expanded by generating all possible 90 degree rotations and horizontal flips for each image in each data partition. Output point annotations were converted to cones centered on the output with a radius of 4 pixels following.^30^ Input output pairs were generated by selecting random 256×256 crops of each input image, then further cropping the output image to match the output detector size (256×256 for Residual U-net, FCRN-A and FCRN-B, 225×225 for Count-ception, 96×96 for U-net).

To establish the number of epochs to train each neural net before saturation, we trained each net for 500,000 epochs and evaluated performance on the test set at the 10, 50, 100, 200, 300, 400, and 500 thousandth epoch using the Adam algorithm with a learning rate of 1e-4. Each neural net had a slightly different loss behavior, but all nets saturated around 100,000 epochs with highest performance on the test data at 50,000 epochs. Each architecture was then trained three times to 100,000 epochs and evaluated at the 10, 25, 50, 75, and 100 thousandth epoch on the test data set. The top three highest performing architectures were then ensembled to maximize train and test score from compositing the individual net segmentations using grid search to weight each net over the range 0.0 to 2.0 inclusive in steps of 0.1. All nets receiver operating characteristic (ROC) and precision and recall were compared on the validation data set, and ranked according to area under the curve (AUC).

To establish how well each network could transfer segmentations between imaging systems, the confocal data set was also split into 80 % training, 10 % test, 10 % validation, and then each fold expanded by generating all possible 90 degree rotations and horizontal flips for each image in each data partition. Neural nets trained only on the inverted data set had poor performance, so each net was additionally trained for 25,000 epochs with evaluations on the test set at the 1, 2, 5, 10, 15 and 25 thousandth epochs using Adam with a learning rate of 1e-5. Weights for the ensemble were recalculated using grid search as described above and net performance was again compared to the validation data set using AUC.

### Cell correspondence algorithm

To detect cells in images of arbitrary size, each neural net was convolved with a zero-padded image to create a final output probability mask with the same size as the original image, with a convolution stride of 1, excluding 5 pixels around each border. To convert individual neural net predictions to cell point detections, the peak detections of each cell center were segmented using non-local maximum suppression with a minimal activation level of 0.1 and a minimal distance of 3 pixels.

Cell correspondence was found by greedily pairing the closest detected cell center in each frame to the next, taking the closest match in cases of multiple linkage, with a maximal link distance of 8 um. Since on average 5% of cells were not detected each frame, nearest neighbor linkage resulted in many short track fragments with single frame breaks that impeded long term cell tracking. To link discontinuous fragments, a dense mesh was imposed on colonies in space using Delaunay triangulation, and then holes in the mesh were detected by finding points connected to at least 3 other neighbor points in one frame, but missing in the next, then imputing their position using the average motion of the neighborhood. Finally, track fragments shorter than 15 minutes (3-5 frames) were removed from the data set as short fragments were found to not correspond to cells.

### Track evaluation

To calculate individual track metrics, each individual track’s x and y coordinates were first interpolated in time by a factor of 4 before smoothing using a rolling average with a filter width of 5 samples. Smoothed tracks were then used to calculate track length, total cell displacement, and velocity. Track persistence was calculated by analyzing the change in direction of travel at each step. A track segment was considered instantaneously persistent if the velocity was greater than 0.91 *μm/min* and did not turn more than 3 degrees/min.

### Spatial metrics

Whole colony metrics were calculated using a Delaunay triangulation after removing links longer than 50 *μ*m (5 cell diameters away). The largest completely connected region was selected as the colony segmentation, and the perimeter and area of the whole region were calculated. To calculate estimates of density at each track point, the area of each triangle surrounding the point was calculated and the density estimated as the inverse of the average of those areas.

To map all colonies to a uniform coordinate system, the colony perimeter was projected onto the unit circle by calculating the angular position of each perimeter point, then using the distance from that perimeter point to the center as the local radius, and finally by gridding these radii onto a radially uniform 500 point grid. All interior points were then projected onto the unit circle by normalizing each point’s radius with the average perimeter radius at the two nearest angular bins.

### Statistical analysis of colony behavior

Average colony spatial behavior was assessed by dividing warped colonies into three annular rings of equal area: the center, a middle ring, and the periphery. Average density, velocity, and persistence were calculated for each bin averaging over all time and over each colony in the experimental group. All possible comparisons for each group and bin were performed using two-sided t-tests with the Bonferroni-Holm correction for multiple comparisons with significance was assessed at p < 0.05. Additionally, 95 % confidence intervals around the mean were calculated using 1000 iterations of bootstrap sampling. Effect size was assessed using Cohen’s d using pooled standard deviations as measured using the maximum likelihood estimator.

---

**Figure S1.**
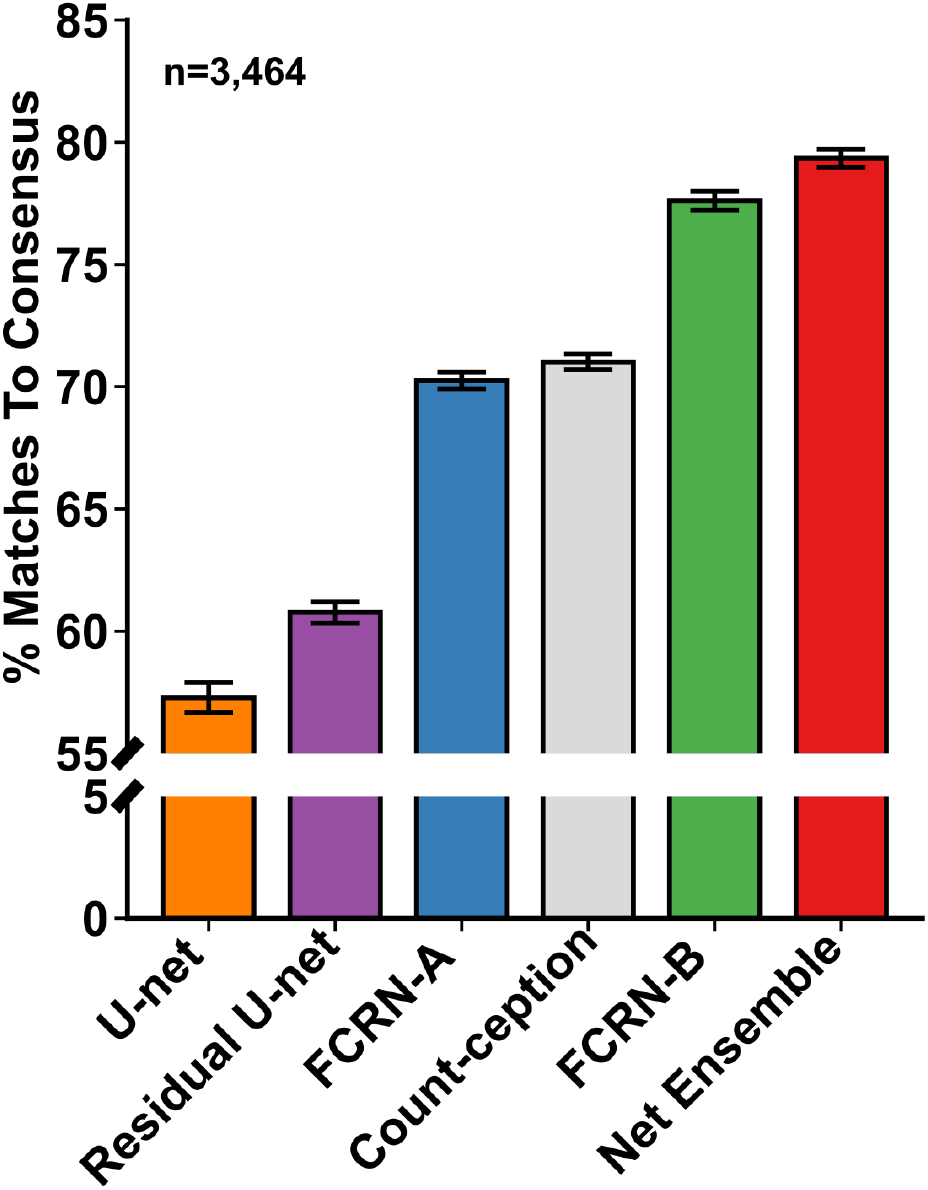
Neural Net Ensemble has Highest Individual Rater Reliability. Individual rater reliability for each neural net architecture compared to human annotated dataset.

**Figure S2.**
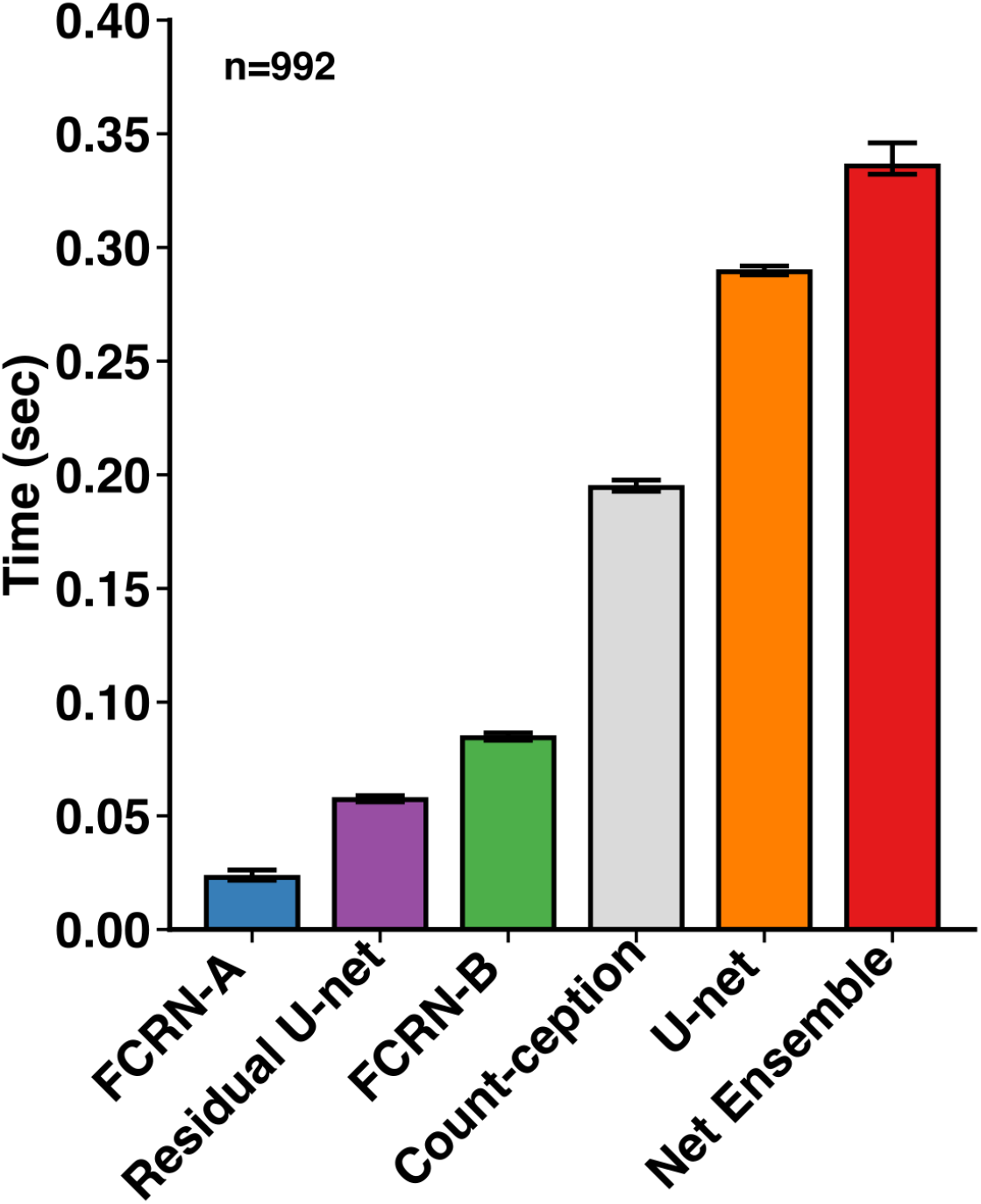
Neural Net Ensemble Segmentation Time Comparable to the Slowest Neural Net. Average time to segment one image for each neural net architecture.

**Figure S3.**
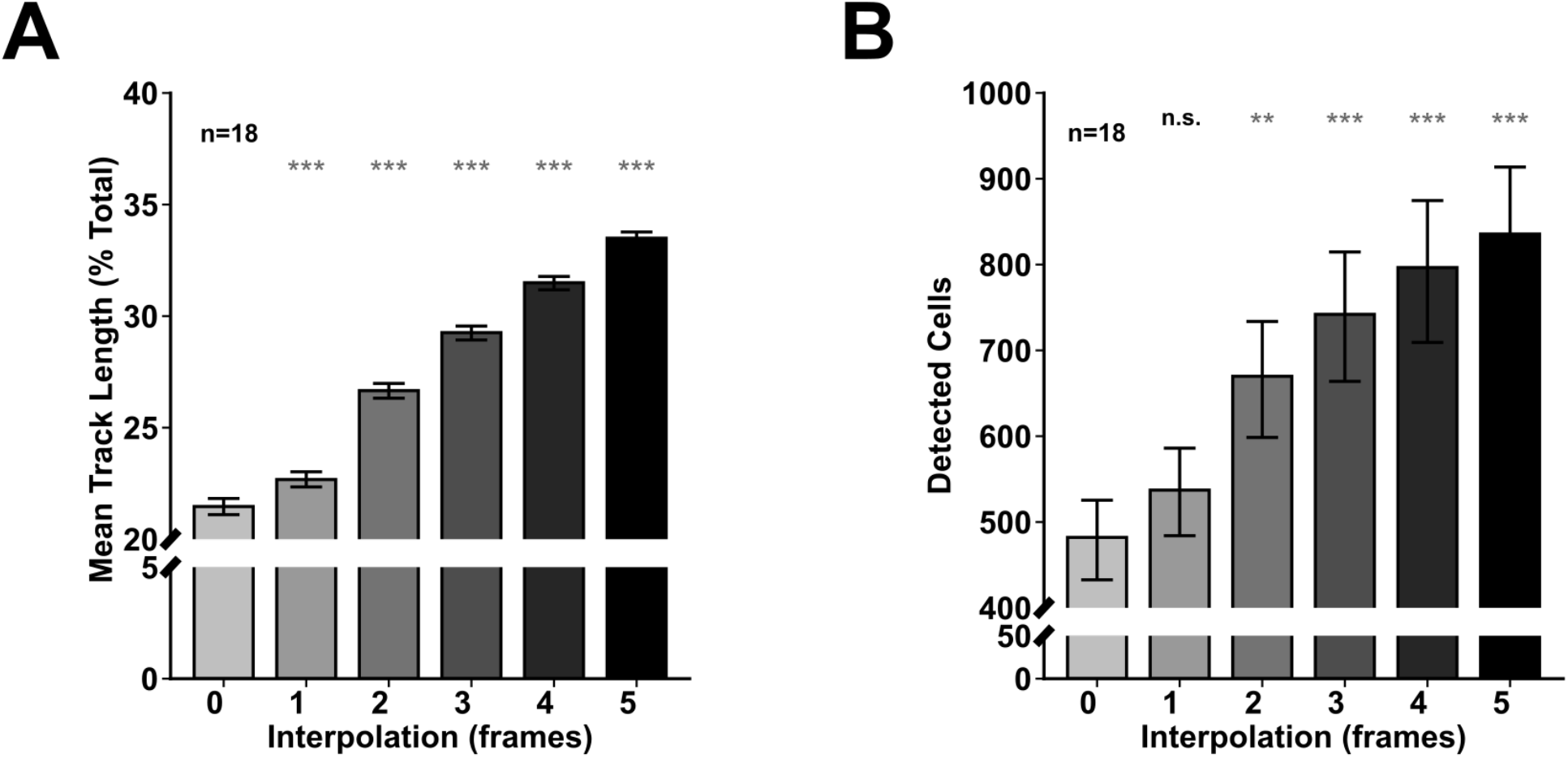
Track Interpolation Across Frames Increases Track Length and Detected Cell Number. A. Mean track length increases with increasing number of interpolated frames (** p < 0.01, *** p < 0.001). B. Mean cell number increases with increasing number of interpolated frames (** p < 0.01, *** p < 0.001).

**Figure S4.**
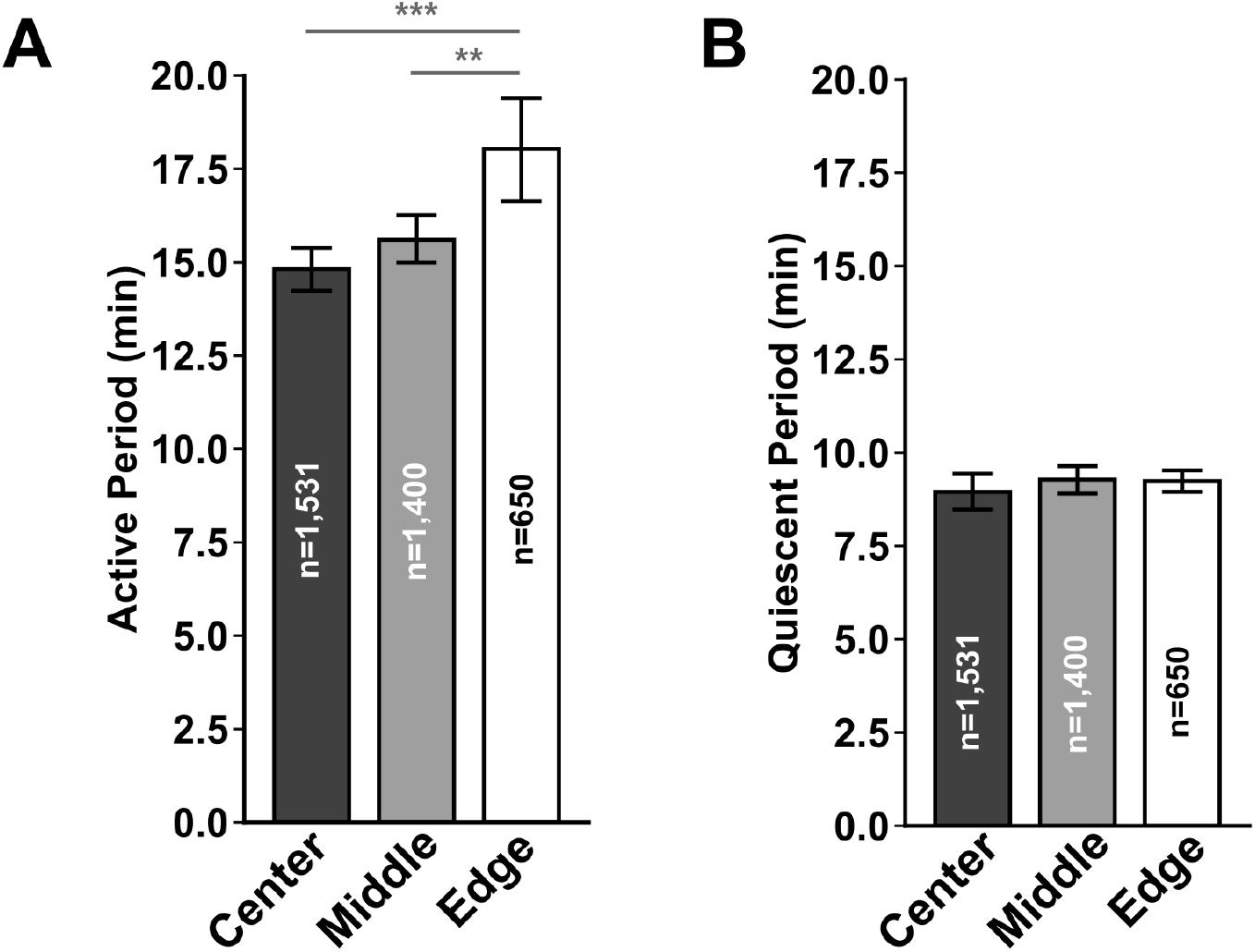
Active Migration Time is Higher at the Colony Edge. A. Average period of active migration is higher at the edge than the center (** p < 0.01, *** p < 0.001). B. Average quiescent period between migrations is not different between the edge and center of colonies

**Figure S5.**
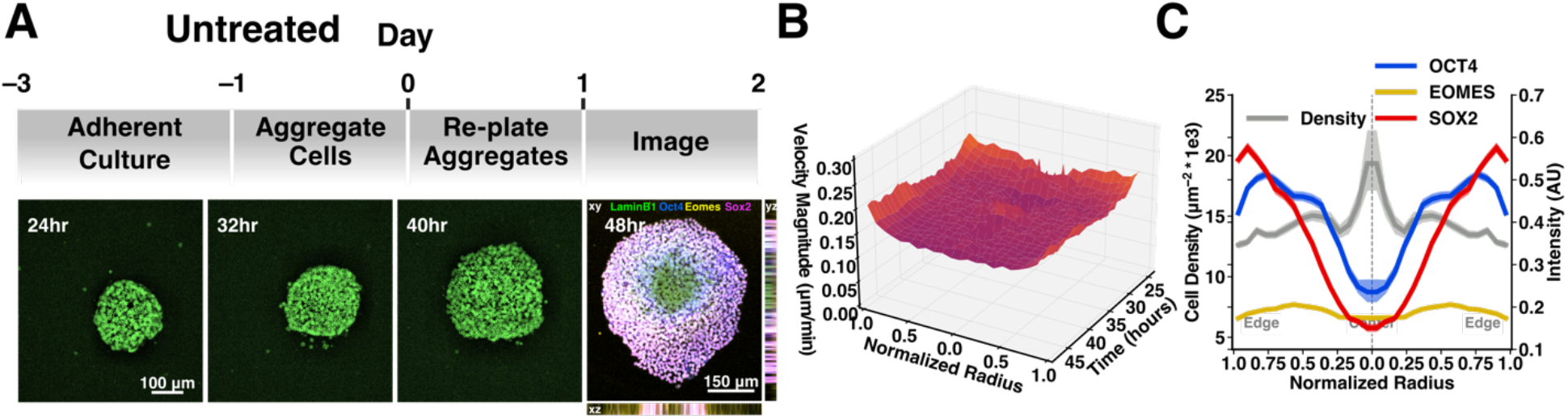
Untreated Colonies Exhibit Uniform Dynamic Behaviors over 24 Hours. A. Treatment timeline and example time course of untreated colonies at 24, 32, and 40 HPR with fixed and stained image of the same colony at 48 HPR. ii. Surface plot of temporal evolution of average instantaneous cell velocity over untreated colonies projected on to the unit circle (n=12 colonies). iii. OCT4, SOX2, and EOMES expression profiles and average cell density profile at 48 HPR projected onto the unit circle in untreated colonies (n=12 colonies).

**Figure S6.**
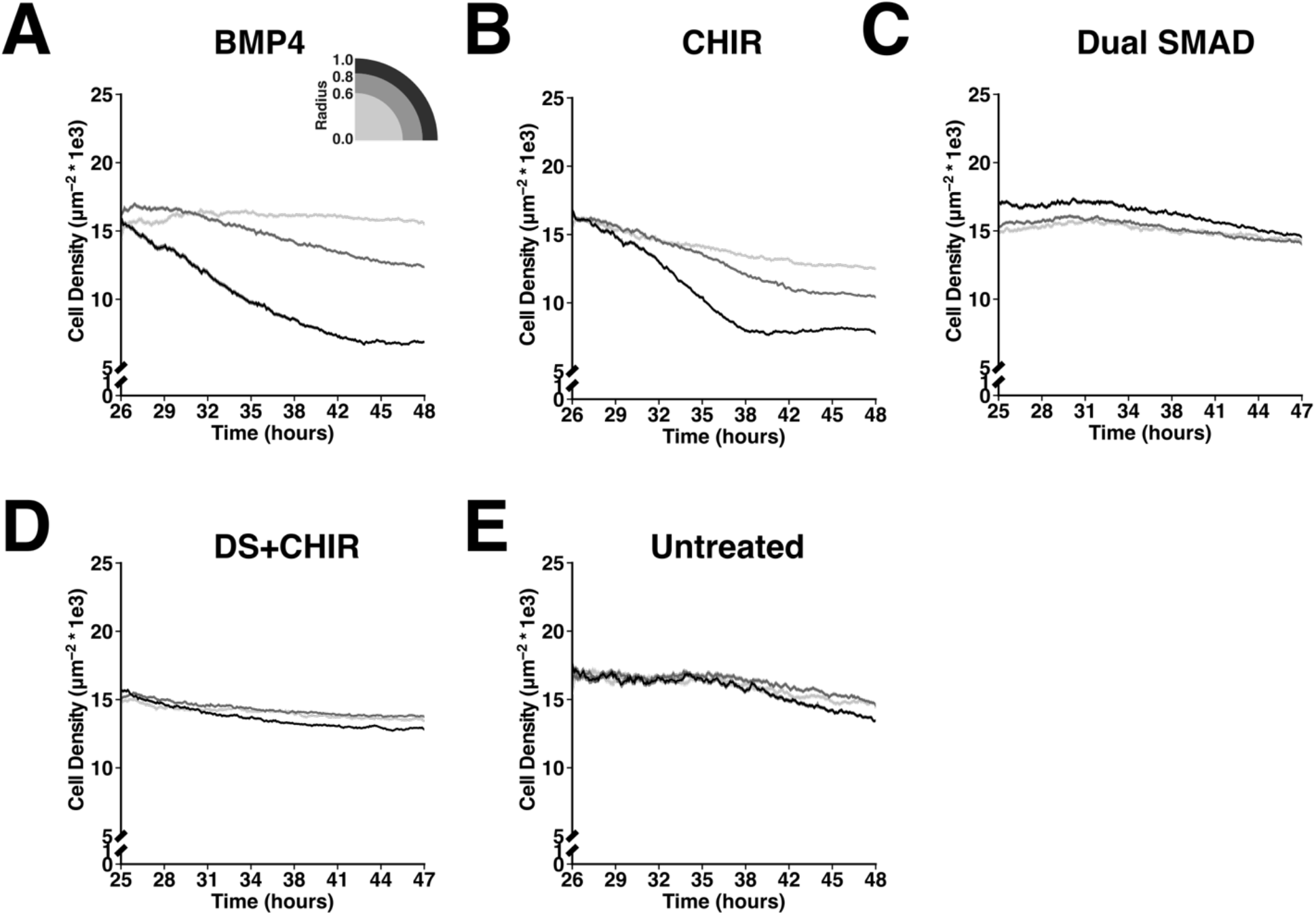
Cell Density Bifurcates in Response to Morphogen Treatment. Temporal evolution of average cell density stratified by colony region (n=16 colonies/condition, n=12 untreated) in A. BMP4 treated colonies, B. CHIR treated colonies, C. Dual SMAD treated colonies, D. Dual SMAD colonies pre-treated with CHIR, and E. untreated colonies.

**Figure S7.**
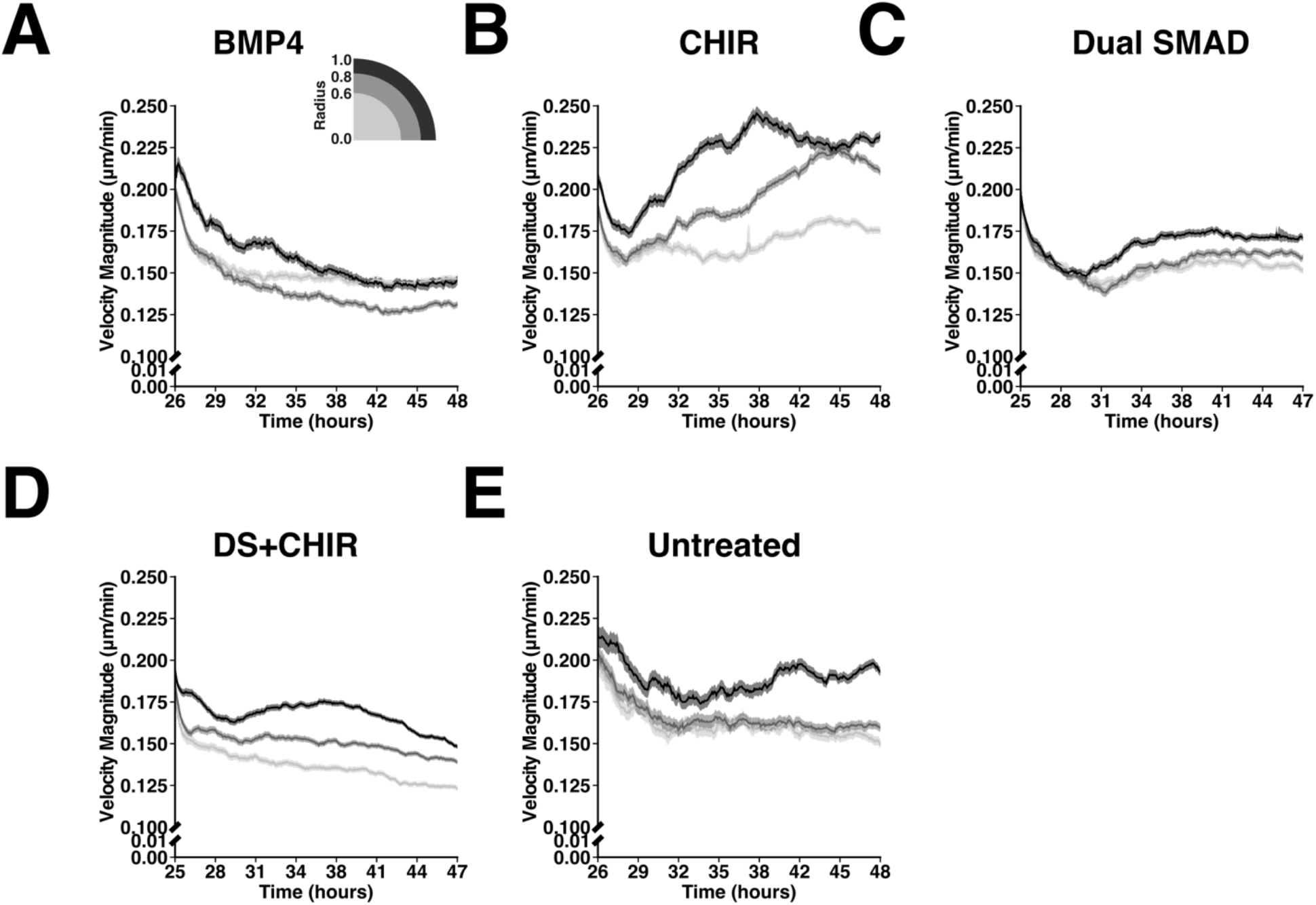
Cell Velocity Magnitude Responds Characteristically to Morphogen Treatment. Temporal evolution of average cell velocity magnitude stratified by colony region (n=16 colonies/condition, n=12 untreated) in A. BMP4 treated colonies, B. CHIR treated colonies, C. Dual SMAD treated colonies, D. Dual SMAD colonies pre-treated with CHIR, and E. untreated colonies.

## Supplementary Tables

**Table S1.** Antibodies

| Gene Target (Symbol) | Species | Company (cat. #) | Dilution |
| --- | --- | --- | --- |
| SRY-box 2 (SOX2) | rabbit | Invitrogen (48-1400) | 1:400 |
| Octamer-binding transcription factor 4 (OCT4) | goat | Santa Cruz Biotechnology (SC-8629) | 1:400 |
| Eomesodermin (EOMES) | mouse | R&D Systems (MAB6166) | 1:400 |

## Supplementary Materials

Figure S1: Neural Net Ensemble has Highest Individual Rater Reliability

Figure S2: Neural Net Ensemble Segmentation Time Comparable to the Slowest Neural Net

Figure S3: Track Interpolation Across Frames Increases Track Length and Detected Cell Number

Figure S4: Active Migration Time is Higher at the Colony Edge

Figure S5: Untreated Colonies Exhibit Uniform Dynamic Behaviors over 24 Hours

Figure S6 - Cell Density Bifurcates in Response to Morphogen Treatment

Figure S7 - Cell Velocity Magnitude Responds Characteristically to Morphogen Treatment

Movie S1: Migration patterns in regions of pluripotent colonies

Movie S2: Migration in BMP4 treated colonies

Movie S3: Migration in CHIR treated colonies

Movie S4: Migration in Dual SMAD inhibition colonies

Movie S5: Migration in CHIR, Dual SMAD inhibition co-treated colonies

Movie S6: Migration in untreated colonies

Table S1: List of antibodies

## Acknowledgements

We would like to thank our human annotators for their hard work segmenting hiPSC colony images. Additionally, we would like to acknowledge the Gladstone Stem Cell Core, the Gladstone Light Microscopy and Histology Core, and the Gladstone Graphics team for their invaluable technical support. Finally, we would like to specifically thank Dr. Kathrine Pollard for her analytical advice and expertise.

## Author contributions

DAJ, ARGL, and TCM conceptualized and designed the experiments. DAJ and ARGL performed the pluripotent cell culture and stem cell differentiations. DAJ performed the data acquisition, with advice from ARGL. Neural network implementation, training, and evaluation were performed by DAJ. The image analysis pipeline was designed, written, and executed by DAJ. DAJ wrote the original manuscript with input from all co-authors. DAJ prepared the figures with input from all co-authors.

## References

1. Sulston, J. E., Schierenberg, E., White, J. G. & Thomson, J. N. The embryonic cell lineage of the nematode Caenorhabditis elegans. Dev. Biol. 100, 64–119 (1983).

2. Chhetri, R. K. et al. Whole-animal functional and developmental imaging with isotropic spatial resolution. Nat. Methods 12, 1171–1178 (2015).

3. Peng, G. et al. Spatial Transcriptome for the Molecular Annotation of Lineage Fates and Cell Identity in Mid-gastrula Mouse Embryo. Dev. Cell 36, 681–697 (2016).

4. Deglincerti, A. et al. Self-organization of the in vitro attached human embryo. Nature 533, 251–254 (2016).

5. Shahbazi, M. N. et al. Self-organization of the human embryo in the absence of maternal tissues. Nat. Cell Biol. 18, 700–708 (2016).

6. Bao, Z. et al. Automated cell lineage tracing in Caenorhabditis elegans. Proc. Natl. Acad. Sci. U. S. A. 103, 2707–2712 (2006).

7. Cai, D., Cohen, K. B., Luo, T., Lichtman, J. W. & Sanes, J. R. Improved tools for the Brainbow toolbox. Nat. Methods 10, 540–547 (2013).

8. Henner, A., Ventura, P. B., Jiang, Y. & Zong, H. MADM-ML, a Mouse Genetic Mosaic System with Increased Clonal Efficiency. PLoS ONE 8, e77672 (2013).

9. Lou, X., Kang, M., Xenopoulos, P., Muñoz-Descalzo, S. & Hadjantonakis, A.-K. A Rapid and Efficient 2D/3D Nuclear Segmentation Method for Analysis of Early Mouse Embryo and Stem Cell Image Data. Stem Cell Rep. 2, 382–397 (2014).

10. Stegmaier, J. et al. Real-Time Three-Dimensional Cell Segmentation in Large-Scale Microscopy Data of Developing Embryos. Dev. Cell 36, 225–240 (2016).

11. Maiuri, P. et al. The first World Cell Race. Curr. Biol. 22, R673–R675 (2012).

12. Libby, A. R. et al. Spatiotemporal mosaic self-patterning of pluripotent stem cells using CRISPR interference. eLife 7, (2018).

13. Turing, A. M. The chemical basis of morphogenesis. Philos. Trans. R. Soc. Lond. B Biol. Sci. 237, 37–72 (1952).

14. Warmflash, A., Sorre, B., Etoc, F., Siggia, E. D. & Brivanlou, A. H. A method to recapitulate early embryonic spatial patterning in human embryonic stem cells. Nat. Methods 11, 847–854 (2014).

15. Hookway, T. A., Butts, J. C., Lee, E., Tang, H. & McDevitt, T. C. Aggregate formation and suspension culture of human pluripotent stem cells and differentiated progeny. Methods 101, 11–20 (2016).

16. White, D. E., Kinney, M. A., McDevitt, T. C. & Kemp, M. L. Spatial Pattern Dynamics of 3D Stem Cell Loss of Pluripotency via Rules-Based Computational Modeling. PLoS Comput. Biol. 9, e1002952 (2013).

17. Novkovic, M. et al. Topological Small-World Organization of the Fibroblastic Reticular Cell Network Determines Lymph Node Functionality. PLOS Biol. 14, e1002515 (2016).

18. Malmersjo, S. et al. Neural progenitors organize in small-world networks to promote cell proliferation. Proc. Natl. Acad. Sci. 110, E1524–E1532 (2013).

19. Barabási, A.-L., Albert, R. & Jeong, H. Mean-field theory for scale-free random networks. Phys. Stat. Mech. Its Appl. 272, 173–187 (1999).

20. Martinez Arias, A. & Steventon, B. On the nature and function of organizers. Development 145, dev159525 (2018).

21. Shahbazi, M. N. & Zernicka-Goetz, M. Deconstructing and reconstructing the mouse and human early embryo. Nat. Cell Biol. 20, 878–887 (2018).

22. LeCun, Y., Bengio, Y. & Hinton, G. Deep learning. Nature 521, 436–444 (2015).

23. Moen, E. et al. Deep learning for cellular image analysis. Nat. Methods 16, 1233–1246 (2019).

24. Xie, W., Noble, J. A. & Zisserman, A. Microscopy cell counting and detection with fully convolutional regression networks. Comput. Methods Biomech. Biomed. Eng. Imaging Vis. 1–10 (2016) doi:10.1080/21681163.2016.1149104.

25. Su, H. et al. Robust Cell Detection and Segmentation in Histopathological Images Using Sparse Reconstruction and Stacked Denoising Autoencoders. in Medical Image Computing and Computer-Assisted Intervention – MICCAI2015 (eds. Navab, N., Hornegger, J., Wells, W. M. & Frangi, A. F.) vol. 9351 383–390 (Springer International Publishing, 2015).

26. Falk, T. et al. U-Net: deep learning for cell counting, detection, and morphometry. Nat. Methods 16, 67–70 (2019).

27. Xie, Y. et al. Efficient and robust cell detection: A structured regression approach. Med. Image Anal. 44, 245–254 (2018).

28. Ronneberger, O., Fischer, P. & Brox, T. U-net: Convolutional networks for biomedical image segmentation. in International Conference on Medical Image Computing and Computer-Assisted Intervention 234–241 (Springer, 2015).

29. Szegedy, C. et al. Going deeper with convolutions. in 2015 IEEE Conference on Computer Vision and Pattern Recognition (CVPR) 1–9 (2015).

30. Cohen, J. P., Lo, H. Z. & Bengio, Y. Count-ception: Counting by Fully Convolutional Redundant Counting. ArXiv Prepr. ArXiv170308710 (2017).

31. Caicedo, J. C. et al. Nucleus segmentation across imaging experiments: the 2018 Data Science Bowl. Nat. Methods 16, 1247–1253 (2019).

32. Ulman, V. et al. An objective comparison of cell-tracking algorithms. Nat. Methods (2017) doi:10.1038/nmeth.4473.

33. Darnton, N. C., Turner, L., Rojevsky, S. & Berg, H. C. On Torque and Tumbling in Swimming Escherichia coli. J. Bacteriol. 189, 1756–1764 (2007).

34. Devreotes, P. & Janetopoulos, C. Eukaryotic Chemotaxis: Distinctions between Directional Sensing and Polarization. J. Biol. Chem. 278, 20445–20448 (2003).

35. Pegoraro, A. F., Fredberg, J. J. & Park, J.-A. Problems in biology with many scales of length: Cell–cell adhesion and cell jamming in collective cellular migration. Exp. Cell Res. 343, 54–59 (2016).

36. Cui, C., Yang, X., Chuai, M., Glazier, J. A. & Weijer, C. J. Analysis of tissue flow patterns during primitive streak formation in the chick embryo. Dev. Biol. 284, 37–47 (2005).

37. Lian, X. et al. Directed cardiomyocyte differentiation from human pluripotent stem cells by modulating Wnt/β-catenin signaling under fully defined conditions. Nat. Protoc. 8, 162–175 (2013).

38. Przybyla, L., Lakins, J. N. & Weaver, V. M. Tissue Mechanics Orchestrate Wnt-Dependent Human Embryonic Stem Cell Differentiation. Cell Stem Cell 19, 462–475 (2016).

39. Libby, A. R. G. et al. Automated Design of Pluripotent Stem Cell Self-Organization. Cell Syst. 9, 483–495.e10 (2019).

40. Glen, C. M., McDevitt, T. C. & Kemp, M. L. Dynamic intercellular transport modulates the spatial patterning of differentiation during early neural commitment. Nat. Commun. 9, (2018).

41. Chambers, S. M. et al. Highly efficient neural conversion of human ES and iPS cells by dual inhibition of SMAD signaling. Nat. Biotechnol. 27, 275–280 (2009).

42. Tewary, M. et al. A stepwise model of reaction-diffusion and positional information governs self-organized human peri-gastrulation-like patterning. Development 144, 4298–4312 (2017).

43. Libby, A. et al. Elongation of Caudalized Human Organoids Mimics Neural Tube Development. http://biorxiv.org/lookup/doi/10.1101/2020.03.05.979732 (2020) doi:10.1101/2020.03.05.979732.

44. Mohammed, H. et al. Single-Cell Landscape of Transcriptional Heterogeneity and Cell Fate Decisions during Mouse Early Gastrulation. Cell Rep. 20, 1215–1228 (2017).

45. Ludwig, T. E. et al. Feeder-independent culture of human embryonic stem cells. Nat. Methods 3, 637–646 (2006).

46. Watanabe, K. et al. A ROCK inhibitor permits survival of dissociated human embryonic stem cells. Nat. Biotechnol. 25, 681–686 (2007).

